# Structural and biochemical characterization of bifunctional XynA

**DOI:** 10.1101/2020.10.20.348094

**Authors:** Wei Xie, Qi Yu, Yun Liu, Ruoting Cao, Ruiqing Zhang, Sidi Wang, Ruoting Zhan, Zhongqiu Liu, Kui Wang, Caiyan Wang

## Abstract

Xylan and cellulose are the two major constituents in numerous types of lignocellulosic biomass, representing a promising resource for biofuels and other biobased industries. The efficient degradation of lignocellulose requires the synergistic actions of cellulase and xylanase. Thus, bifunctional enzyme incorporated xylanase/cellulase activity has attracted considerable attention since it has great cost savings potential. Recently, a novel GH10 family enzyme XynA identified from *Bacillus* sp. is found to degrade both cellulose and xylan. To understand its molecular catalytic mechanism, here we first solve the crystal structure of XynA at 2.3 Å. XynA is characterized with a classic (α/β)8 TIM-barrel fold (GH10 domain) flanked by the flexible N-terminal domain and C-terminal domain. Circular dichroism, protein thermal shift and enzyme activity assays reveal that conserved residues Glu182 and Glu280 are both important for catalytic activities of XynA, which is verified by the crystal structure of XynA with E182A/E280A double mutant. Molecular docking studies of XynA with xylohexaose and cellohexaose as well as site-directed mutagenesis and enzyme activity assay demonstrat that Gln250 and His252 are indispensible to cellulase and bifunctional activity, separately. These results elucidate the structural and biochemical features of XynA, providing clues for further modification of XynA for industrial application.

## Introduction

With the continuous consumption of oil resources and the development of the global economy, developing cleaner and more economical energy is of utmost importance (1). Biomass energy is such a kind of renewable energy (2, 3) and biomass has diverse and abundant sources including forestry byproducts and crop straws. During production, the plant cell walls must be depolymerized, which requires synergistic action of cellulase and xylanase to degrade the ingredients of plant cell walls including cellulose, hemicellulose, pectin, and lignin (4–7). In addition to biofuel production, xylanase is also applied in various industries, such as the food industry, feed industry, paper industry, and brewing industry (8–11).

The thermostable glycoside hydrolase (GH) family with both cellulase and xylanase activity is more advantageous in degrading lignocellulose biomass, and thus has great development potential for industrial application. Among the GH family, the GH10 family and GH11 family contain a large number of xylanases (12). Compared with GH11 xylanases, GH10 xylanase has an extensive substrate scope (13–15). Furthermore, the GH10 xylanase can also degrade other polysaccharides, such as konjac glucomannan and tamarind xyloglucan (16). The GH10 domain of GH10 family xylanases is responsible for catalytic activity. The region outside the GH10 domain (N-terminal or C-terminal) determines the specificity for different substrates (17). Because the binding ability of xylanase and the substrate is relatively weak, it is difficult to detect their binding, such as the GH10A xylanase from the Arctic mid-ocean ridge ventilation system. The xylanase GH10A is capable of degrading wheat arabinoxylan (WAX) and tamarind xyloglucan and it is difficult to detect the binding ability (16).

In a previous study, a novel GH10 xylanase termed XynA was identified from *Bacillus* sp. (18). XynA could function at a wide range of pH (maintains more than 60% activity at pH 5.0-7.5) and temperature (maintains more than 65% activity at 45 °C to 80 °C) (18). Different from other GH10 enzymes, XynA is a special bifunctional xylanase/cellulase enzyme that can degrade both xylan substrates and a variety of cellulose substrates, including cellobiose, cellohexaose, carboxymethyl cellulose, filter paper, Avicel (microcrystalline cellulose), *p*-nitrobenzene-cellobioside and *p*-nitrobenzene-glucopyranoside (18). In addition, XynA hydrolyzed corn stover pretreated with cellulase, showing an obvious synergistic effect (18). These unparalleled characteristics make XynA become an attractive candidate in biotechnology applications, such as bioenergy production and pulp processing (19–21). However, the xyanase activity of XynA is much stronger than its cellulase activity, which may raise some problems during application, for instance, the insufficient degradation of substrates and the quality control.

To provide useful guidelines for design of XynA for future industrial application, we started from structure to investigate the catalytic mechanisms of the xyanase and cellulase activity of XynA. We report the crystal structure of bifunctional enzyme, XynA at 2.3 Å firstly. Then from structural clues, we designed function assays and revealed that Glu182 and Glu280 are conserved and crucial for XynA. Gln250 is critical to cellulase activity of XynA. His252 is critical to bifunctional activity of XynA. The N-terminal and C-terminal of XynA interact with the GH10 domain through hydrogen bonds. The interaction between N-terminal and GH10 domains as well as C-terminal and GH10 domains and their conformational changes affect the activity and stability of XynA, indicating the importance of N-terminus and C-terminus.

## Results

### The overall structure of XynA

The full-length XynA protein was expressed and purified with removal of nucleic acids by anion exchange column QHP. XynA contains 408 residues and the molecular weight of the tagged full-length protein was 49.8 kDa with an isoelectric point of 6.64. It was identified as one monomer in the solution by gel filtration (Figure 1A). We found that xylanase 10A (SlXyn10A, PDB code 1V0L) from *S. lividans* has the highest sequence similarity (30%) with XynA by using Phyre2 server (22, 23). We used this structure as a search model to solve the crystal structure of XynA by molecular replacement method.

**Figure 1.**
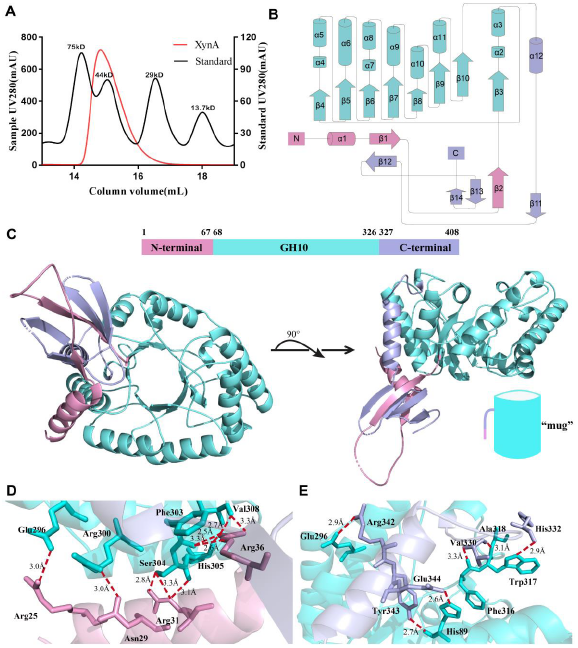
XynA protein and its structure. (A) Gel-filtration analysis showing XynA as monomer in solution. The red and black lines represent the UV absorption of XynA and the standard proteins at 280 nM respectively. (B) The topological organization of XynA. The GH10 catalytic domain, the N-terminal domain and C-terminal domain are colored in cyan, pink and purple, respectively. (C) Overall structure of XynA is like a shape of “mug”. The color is assigned as shown in front. (D) The hydrogen bonding interaction between the N-terminal and GH10 domains. The red dashed line indicates hydrogen bonding interaction. (E) The hydrogen bonding interaction between the C-terminal and GH10 domains.

The determined 2.3 Å crystal structure of XynA (residues 22-393) with a *R*_free_ of 0.245 (Table 1). The space group is *P4122*, and one molecule was present in the asymmetric unit. In the refined model, residues Leu52, Ala53, Asp359, Ala360, Asn361, Gly378, and Glu379 are missing due to disordered configuration. The XynA structure is composed of 12 α-helices and 14 β-sheets, among which α2-α11 helices and β3-β10 sheets form a classical (α/β)_8_ TIM-barrel fold (Figure 1B). Overall, the mug-shaped XynA is comprised of an N-terminal domain (residues 1-67), GH10 domain (residues 68-326) and C-terminal domain (residues 327-408) (Figure 1C). The body of “mug” is GH10 domain, which has 3 α-helices separated by loop regions, named α2, α4, and α7 (Figure 1B). These three α-helices are not affecting the overall conformation of classical (α/β)_8_ TIM-barrel fold. The N-terminal and C-terminal form the handle of this “mug” together. We find that the N-terminal residues Arg25, Asn29, Arg31, and Arg36 establish abundant hydrogen bond interactions with the residues Glu296, Arg300, Phe303, Ser304, His305, and Val308 at the body GH10 domain (Figure 1D). Similarly, the C-terminal residues Val330, His332, Arg342, Tyr343 and Glu344 form extensive hydrogen bond interactions with residues His89, Glu296, Phe316, Trp317, and Ala318 of the GH10 domain (Figure 1E). The residues participated in hydrogen bond interactions are all located in α-helices and loops. The difference between XynA and other GH10 families exists in the N-terminal and C-terminal domain. The C-terminal residues (Glu338-Glu349) form α-helix_12_, which makes the classical (α/β)_8_ TIM-barrel fold more complete. The N-terminal (α1, β1 and β2) and C-terminal domain (α12, β11-β14) is partially crossover, and there are two α-helices and six β-sheets that are adjacent to the TIM barrel.

**Table 1.** Data collection and refinement statistics.

|  | XynA WT | XynA E182A/E280A |
| --- | --- | --- |
| **Data collection** |  |  |
| Wavelength (Å) | SSRF-BL19U1 | SSRF-BL19U1 |
| Space group | *P*4122 | *P*4122 |
| a, b, c (Å) | 109.842, 109.842, 85.57 | 110.193, 110.193, 84.983 |
| $\alpha, \beta, \gamma$ (°) | 90.00, 90.00, 90.00 | 90.00, 90.00, 90.00 |
| Resolution (Å) | 15.29-2.345 (2.429-2.345)<sup>a</sup> | 49.28-2.894 (2.997- 2.894)<sup>a</sup> |
| $R_{\text{merge}}^b$ | 0.158 (1.074) | 0.108(0.550) |
| $I/\sigma(I)$ | 30(2.6) | 34(5.83) |
| Completeness (%) | 99.35 (98.73) | 99.56 (96.27) |
| Redundancy | 25.3(23.6) | 25.5(27.1) |
| **Refinement** |  |  |
| Resolution (Å) | 15.29-2.345 | 49.28-2.894 |
| No. reflections | 22400 (2182) | 12123 (1135) |
| $R_{\text{work}}^c/R_{\text{free}}^d$ | 0.2213/0.2446 | 0.2378/0.2690 |
| No. atoms | 3041 | 2884 |
| Protein | 2972 | 2856 |
| Ligand | 2 | 2 |
| Water molecules | 67 | 26 |
| B-factors (Å<sup>2</sup>) |  |  |
| Protein | 55.49 | 63.42 |
| Ligand | 51.66 | 49.77 |
| Water molecules | 56.34 | 47.08 |
| Bond lengths (Å) | 0.009 | 0.014 |
| Bond angles (°) | 1.08 | 1.53 |
| Ramachandran favored (%) | 95.51 | 96.21 |
| allowed | 3.93 | 3.21 |
| Outliers (%) | 0.56 | 0.58 |
| PDB code | 7D88 | 7D89 |

<sup>a</sup>Values in parentheses are for the highest-resolution shell. <sup>b</sup> $R_{\text{merge}} = \sum (I - \langle I \rangle) / \sum I$ , where $I$ is the observed intensity. <sup>c</sup> $R_{\text{work}} = \sum_{\text{hkl}} ||F_o| - |F_c|| / \sum_{\text{hkl}} |F_o|$ , calculated from working data set. <sup>d</sup> $R_{\text{free}}$ is calculated from 5.0% of data randomly chosen and not included in refinement.

### Sequence and structure analyses of XynA with other GH10 xylanase

To find the catalytic residues of XynA, we performed multiple sequence alignments of XynA with five GH10 family enzymes, namely CbXyn10C (*C. bescii*), SlXyn10A (*S. lividans*), BsXynA (*Bacillus sp.*), TmXynB (*T. maritima*) and TsXy1A (*T. saccharolyticum*). Among them, XynA, CbXyn10C and SlXyn10A are xylanases with xylanase and cellulase activity. BsXynA, TmXynB, and TsXy1A are strict xylanase enzyme. We find that the residues in the GH10 domain, such as Phe90, Glu96, Lys100, Gly133 and Trp137 are much conserved (Figure 2A). Though XynA and BsXynA both come from *Bacillus* sp., BsXynA is a traditional GH10 family xylanase that only catalyzes the degradation of xylan (24). The architecture of GH10 domain is similar in XynA and BsXynA, but the subtle sequence difference in fifteen residues indicates that these sites may determine the monofunction or bifunction (Figure 2A). By structural superposition and sequence comparison among CbXyn10C, SlXyn10A, TmXynB, TsXy1A and XynA, we infer that Glu182 and Glu280 are putative catalytic residues of XynA (25–28). Of note, only the GH10 domain is in the other protein structures mentioned above, the N-terminal and C-terminal domains are unique for XynA (Figure 2B). The typical GH10 domain is well conserved in GH10 family, as evidenced by the low RMSD (1.599 Å) of structural superimposition of XynA, TsXy1A and SlXyn10A.

**Figure 2.**
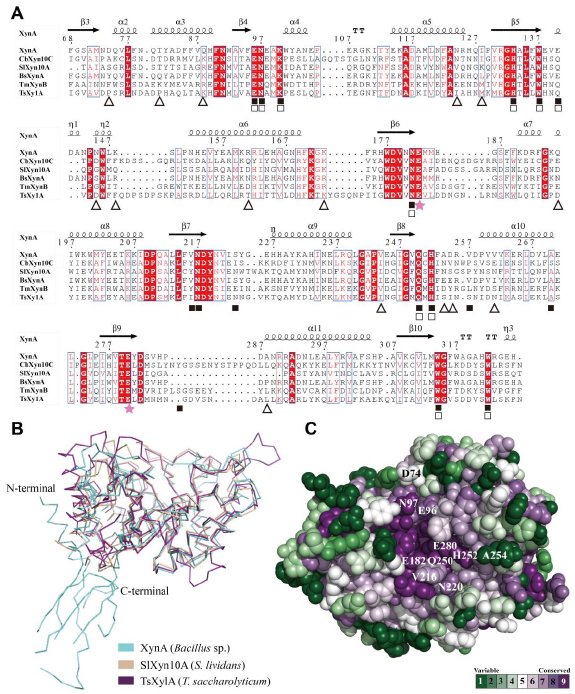
Multiple sequence alignment and structure analyses of XynA. (A) Multiple sequence alignment of XynA (QCO69162) with CbXyn10C (ACM60945) from *C. bescii*; SlXyn10A (AAC26525) from *S. lividans*; BsXynA (AHH02587) from *Bacillus* sp.; TmXynB (AAD35164) from *T. maritima*; and TsXy1A (AFK86466) from *T. saccharolyticum.* The red background color represents strictly conserved residues, and the red font represents the substitution of conserved residues. The pink five-pointed star indicates the two key catalytic residues of XynA, and the original secondary structure of XynA is shown above the sequence alignment. The black squares represent the residues that interact with the substrate in CbXyn10C, and the white squares represent the residues that interact with the substrate in TsXy1A. The triangles indicate the residues of XynA and BsXynA that differ in the GH10 domain. (B) Structural superposition of XynA, SlXyn10A (PDB code 1V0L), and TsXy1A (PDB code 3W25). (C) The spatial position and conservation of residues in space fill model of XynA that may interact with the substrate. The green to purple gradient indicates the degree of conservation. The annotated residues are the mutation sites found by molecular docking.

Further analysis indicates that most residues related to monofunctional and bifunctional activity are conserved and overlaped, containing Glu96, Asn97, Lys100, His133 and Trp137. The residues specific to bifunctional activity include Tyr216, Asn217, Ile223, Trp257 and Tyr288. Among the residues in putative substrate binding pocket, only five residues are varied between bifunctional enzymes XynA and CbXyn10C, namely 140, 216. 223, 257 and 288, indicating conservation in sequence and structure (Figure 2C). The centric Glu182 and Glu280 are surrounded by Glu96, Asn97, Lys100, Tyr216 and Asn217, forming a pocket embracing the sugar chains of substrate.

### The xylanase activity of XynA

To explore the catalytic properties of XynA, we first investigated the optimal enzyme concentration and the xylanase activity of XynA. Using 0.5% (w/v) beechwood xylan (BWX) as substrate in 100 mM sodium citrate buffer, pH6.5 at 65°C for10 minutes, as the concentration of XynA protein increased, the percentage of degraded sugar increased until it reached a plateau (Figure 3A). Based on this reaction curve, 500 nM is regarded as the optimal concentration for XynA.

**Figure 3.**
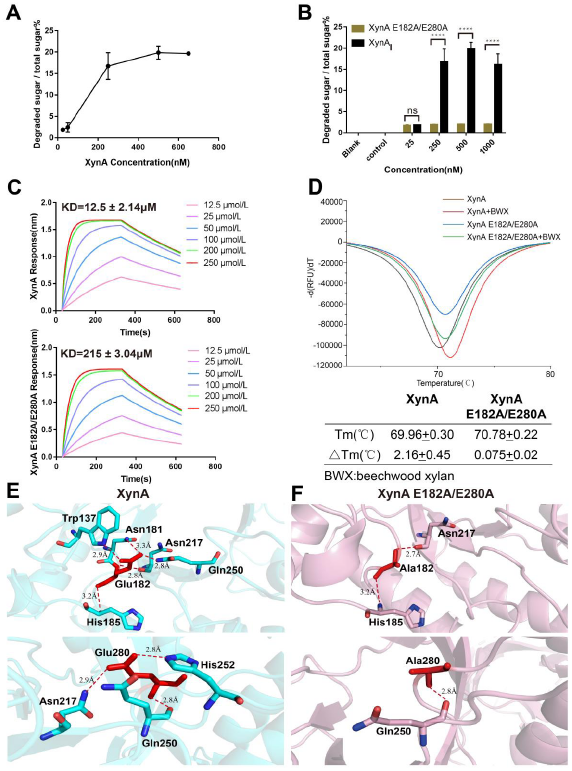
Comparison of XynA with XynA E182A/E280A in terms of binding, catalysis activity and structures. (A) Effect of XynA concentration on its xylanase activity. (B) XynA and XynA E182A/E280A activities for degrading substrates at different concentrations. Each experiment was repeated three times. XynA and XynA E182A/E280A were analyzed for significance, ns means no significant difference, * means P <0.05, ** means P <0.01, *** means P <0.0002, and **** means P <0.0001. (C) XynA binding experiment with BWX; XynA E182A/E280A binding experiment with BWX. Different colored lines in the figure represent different concentrations of BWX. The binding constants are shown in the table. (D) Protein thermal shift assay of XynA and XynA E182A/E280A. The Tm value represents the denaturation or exposure temperature of the hydrophobic residues of the protein. ΔTm indicates the temperature change after adding BWX. (E) Interaction between two residues Glu182 and Glu208 and surrounding residues of XynA. (F) The interaction between the Ala182 and Ala208 residues and the surrounding residues of XynA E182A/E280A. Color cyan and pink indicate the crystal structure of WT and double mutant respectively.

The activity of GH10 family enzymes requires two conserved catalytic residues glutamate, and the mutation of glutamate will completely inactivate the protein (24). To verify the effect of these two glutamic acids on XynA activity, we double mutated Glu182 and Glu280 to alanine and produced the mutant XynA E182A/E280A protein. The xylanase activity of the mutants (XynA E182A/E280A) was determined using the DNS method. The results displayed that WT had the highest activity at a concentration of 500 nM, while the E182A/E280A mutant is inactive, even at an extremely high concentration (1 mM) (Figure 3B).

To analyze the binding mode and interaction patterns of XynA and its substrates, we have tried hard to prepare the crystals of XynA WT, E182A/E280A or E182Q/E280Q with cellobiose, cellohexaose, or xylohexaose complexes, respectively. However, all trials failed, only the apo crystals of WT and mutant XynA were obtained. This may be explained by the weak and difficult detection features for the GH10 family proteins and their substrates according to previous studies (29). However, through Surface Plasmon Resonance (SPR) assay, we detected the binding affinity of BWX to both the WT and E182A/E280A mutant XynA (Figure 3C). The binding constants indicated that the binding ability of the WT to BWX was stronger than that of the E182A/E280A mutant. We found that WT does not bind to oligosaccharides, which is consistent with the fact that it is difficult for oligosaccharides to enter the crystal structure (Figure 3C).

The protein thermal shift assay indicates that XynA is much thermostable, as evidenced by the Tm value (70 °C). Moreover, the Tm value of XynA bound with BWX is higher than that of the apo protein, indicating that substrate binding could stabilize XynA (Figure 3D). However, the Tm value of the mutant before and after BWX bindingis almost unchanged, indicating that substrate binding had no significant effect on the stability of the mutant.

Crystal structure can provide direct hints to study the protein features and properties. To elucidate how E182/E280 affect the enzyme activity of XynA, we determined the crystal structure of XynA E182A/E280A at 2.9 Å. The overall architecture of XynA E182A/E280A is highly similar to the WT XynA, yielding a Cα root-mean-square deviation (RMSD) of 0.218 Å. A major difference observed between the WT and mutant is an angle swing in the N- and the C-terminus. We found that the previously established hydrogen bonds between Glu182 and Trp137, Asn181 and Gln250 as well as Glu280 and with Asn217 and His252 disappeared in the E182A/E280A mutant (Figure 3E and 3F).

### Molecular docking of XynA with xylohexaose and cellohexaose and the mutant activity confirmation

Since the cocrystal structure of XynA complexed with substrates is unsuccessful, to investigate the catalytic mechanism of XynA, we performed molecular docking studies. The sugar chains of xylan and cellulose are too long to suitable for molecular docking, therefore, xylohexaose and cellohexaose are substrates for docking. Viewing the putative binding mode, these two substrates situated in the center of the mouth of the XynA “mug” structure (Figure 4A, Figure 4A 4B). Xylohexaose formed ten hydrogen bonds with surrounding residues in the active pocket (mug mouth), including Asp141, Asn143, Glu96, Lys100, Trp314, Asn220 and Ser223. Cellohexaose formed eleven hydrogen bond interactions with surrounding residues, such as Asp74, Glu96, Glu98, Try102, Glu182, Gln250, His252 and Asn220. We also find that the residues that interact with the substrate in molecular docking are very conservative, and most are located around two catalytic residues (Figure 2C). Based on molecular docking results, residues Gln250 and His252 are speculated with relation to the bifunctionality of XynA. According to the results, Asp74, Glu96, Gln250, His252 and Asn220 formed hydrogen bond with cellohexaose. Based on the classification of residues, we designed mutants D74A, E96A, Q250A, Q250E, H252A and N220E. In light of the residues of CbXyn10C interacting with the substrate, Val216 and Ala254 are potential residues that could interact with the substrate in the XynA sequence (25). We designed mutants N97E, V216E and A254E. We used WAX, BWX and barley β-glucan as substrates, and the activities of these mutants relative to that of the XynA (abbreviated as WT in Figure 4) was determined by the *p*-hydroxybenzoic acid hydrazide (*p*HBAH) method (Figure 4C, 4D, 4E). We found that the four mutants (E182A/E280A, E182Q/E280Q, Q250A, and H252A) were inactive towards three substrates. The mutants E96A and N97E lost their activity towards BWX and were unable to degrade WAX and barley β-glucan by more than 50%. The mutants V216E, N220E, and A254E showed significantly reduced degradation activity on WAX, BWX, and barley β-glucan. The mutant Q250E showed reduced degradation activity on WAX and BWX and no activity against barley β-glucan. The mutant D74A showed significantly weaker degradation activities against WAX and barley β-glucan. E182A/E280A and E182Q/E280Q were completely inactive against these three substrates, further confirming that Glu182 and Glu280 were the key catalytic residue residues.

**Figure 4.**
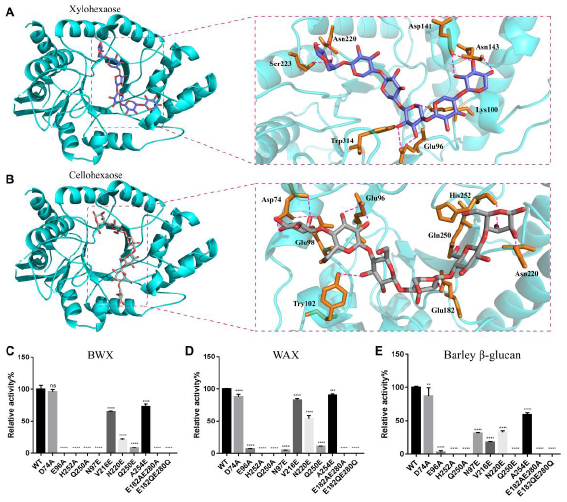
The molecular docking with xylohexaose and cellohexaose and activity analysis of the XynA. (A-B) The docking of XynA with xylohexaose and cellohexaose displayed in a cartoon mode, respectively, and the hydrogen bonds between the XynA with xylohexaose or cellohexaose are partially enlarged. The red dotted line indicates hydrogen bonding interaction. (C-E) Relative activities of the XynA mutants in the hydrolysis of BWX, WAX and barley β-glucan respectively. Each experiment was repeated three times. WT enzyme activity was set to 100%. WT group was used as a control for a significant analysis, * means P <0.05, ** means P <0.01, *** means P <0.0002, and **** means P <0.0001.

## Discussion

The most important features of XynA are its thermal stability, bifunctionality and low homology. It has been reported that its half-lives at 65 °C and 70 °C are 12 hours and 1.5 hours, respectively (18). We can take advantage of the thermal stability of this protein. We tried to purify the protein by heating it in a water bath at 65 °C for 1 hour, which may affect the structure of the protein, so we also used an anion-exchange column QHP for purification. During the crystallization process, we found that the protein purified by the QHP column crystallized quickly, and the crystal morphology was regular, while the protein purified by heating crystallized slowly, and the crystal morphology was irregular. These two kinds of crystallization indicate that the high temperature influences the conformation and stability of the protein, resulting in differences in the crystallization.

In addition to the typical (α/β)_8_ TIM-barrel fold GH10 domain structure, XynA has N-terminal and C-terminal. By homology analysis of the overall structure of XynA, the homology of the “mug handle” formed by the N-terminal and C-terminal parts of XynA is low, the β-sheet conservatism inside the “mug” is high, and the outer part of the α-helix outside the “mug” is not conserved, while the inner part of the α-helix is very conserved (Figure 2C). Structural analysis found that the 39-56 fragments belonging to the N-terminus and 357-367 belonging to the C-terminus are very flexible and the amino acid sequence is not conserved. It has been reported that the N-terminus and C-terminus might affect the thermal stability, activity and substrate specificity of the protein (18, 30–32). The N-terminus and C-terminus are interlaced to form a “mug handle” and they respectively generate hydrogen bond interactions in the GH10 domain (Figure 1D, 1E). Therefore, we think that the changes of the N-terminal and C-terminal will affect the GH10 domain, thereby affecting the activity and stability of XynA. We can consider changing the above two fragments (39-56 and 357-367) to further improve the stability of XynA. We can optimize the activity of XynA from the residues that interact with the GH10 domain. Our analysis showed that the inside of the “mug” is bound to the substrate and degrades the substrate, so the residues are more conserved, while the outside of the “mug” is not bound to the substrate, so the residues change more. We found and verified 2 key residues Glu182 and Glu280 by structural analysis of XynA. In the catalytic process, Glu182 and Glu280 are catalytic nucleophiles/bases and catalytic proton donors, respectively (33). Comparing the properties of WT with those of the double mutant showed the activities, structures and stabilities were all different, proving the importance of Glu182 and Glu280. Glu182 and Glu280 formed hydrogen bonds with surrounding residues could “lock” the substrate to be catalyzed after entering the active pocket. The hydrogen bonds formed by the side chains of Glu182 and Glu280 with surrounding residues account for the majority interaction. Since the residues 182 and 280 are located in the loop area, the reduced interaction also makes this part more flexible, thereby reducing the binding affinity to substrate. This is in consistent with the SPR result. At the same time, WT has the ability to hydrolyze the substrate, while the E182A/E280A has no hydrolysis ability, which also shows that the mutation affects the hydrolysis activity by affecting the binding of the substrate. We attempted to determine the cocrystal structure of the proteins (WT, E182A/E280A, and E182Q/E280Q) with substrates xylohexaose and cellohexaose to reveal the specific catalytic mechanism. However, neither the incubation of the protein with the substrate nor the immersion of the crystal in the substrate successfully afforded cocrystals. From the binding results, we also observed that XynA is weakly bound to the substrate. The substrate is difficult to be fixed in the protein, so it is difficult to obtain the cocrystal structure.

We obtained the simulated structures of XynA and xylohexaose and of XynA and cellohexaose by molecular docking. Cellohexaose has one more -CH_2_OH group than xylohexaose in its molecular structure. Therefore, cellohexaose requires more space than xylohexaose to accommodate the sugar chains. It is found from the molecular docking structure that the XynA active pocket can hold xylohexaose and cellohexaose. This shows that XynA can accommodate sugar chains of different substituents and thus exhibit a broad substrate scope. We indicate that XynA forms multiple hydrogen bonds with xylohexaose and cellohexaose. Compared with the residues that formed hydrogen bonds with xylohexaose and cellohexaose, Asp74, Glu182, Gln250 and His252 are unique residues that form hydrogen bonds between XynA and cellohexaose. According to the molecular docking, Glu96, Lys100, Gln250 and His252 are particularly conserved in the residue sequences of the GH10 family and interact with substrates in CbXyn10C and SlXyn10A.

We find that four mutants, E182A/E280A, E182Q/E280Q, Q250A, and H252A are all inactive towards three substrates. We are not surprised that Glu182 and Glu280 are required for activity, as our results show that they are key catalytic residues. After being replaced with alanine, the activities for the three substrates completely disappeared, which proves that His252 and Gln250 also related to catalysis. Our analysis reveals the conservation of His252 in the GH10 family and the uniformity of its interactions with substrates. The mutant H252A only reduces the activity of the CbXyn10C protein (29). However, XynA H252A loses its activity towards the three substrates, proving that His252 is only an important residue in XynA. For most of the xylanases in the GH10 family, the catalytic residues are Glu (29). But we find that in addition to Glu182 and Glu280 as catalytic residues, XynA also has His252. Gln250, which are also extremely conserved in the residue sequence, and we indicate that Gln250 is the residue that forms hydrogen bonds with both Glu182 and Glu280. In molecular docking, Gln250 also interacts with cellohexaose, so we repute that a hydrogen bonding network is formed around Gln250 to mediate the role of Glu182 and Glu280. The mutation of Gln250 in CbXyn10C decrease the catalytic activity for both xylan and cellulose as substrates (29). The mutant Q250A is inactive against three substrates, while the mutant Q250E still retains activity against xylan substrates. This fully shows that the side chain of Gln250 plays an important role in catalyzing the xylan substrate. The mutant Q250A and Q250E can not hydrolyze the cellulose substrate, indicating that -NH_2_ in the side chain of Gln250 is related to the cellulase activity of XynA. The difference between Gln and Glu is that -NH_2_ in the side chain becomes -OH, and -NH_2_ is more nucleophilic than -OH. We believe that Gln250 is the key residue of XynA with cellulase activity and can be used as a key design element to design mutants of XynA with improved cellulase activity.

The mutants D74A, E96A, N97E, N220E, V216E and A254E all have reduced activity to various extents, proving that these sites all contribute to the activity of the substrate. Homology analysis of the XynA protein structure showed that Gln250 and His252 are very conserved and located between Glu182 and Glu280 in the spatial position (Figure 2C). The four amino acids are very close in space and form multiple hydrogen bonds, proving that Gln250 and His252 play important roles in the activity. We indicate that the residues that interact with Glu182 including Trp137, Asn181, Asn217 and Gln250 and the residues that interact with Glu280 including His133, Asn217, Gln250 and His252 are very conserved in CbXyn10C, and these residues interact with the substrate, fully reflecting the unity of the GH10 family. Finally, we propose the catalytic mechanism of XynA. In the first glycosylation step, E280 acts as a nucleophile (since Glu280 has less interaction with surrounding residues than Glu182, Glu280 is more flexible to generate electron transfer first), attacking the anomeric center to displace glycosides and form glycosylase intermediates. At the same time, Glu182 acts as an acid catalyst and protonates the glycoside oxygen when the bond is broken. In the second deglycosylation step, the glycosylase is hydrolyzed by water, and Glu182 acts as a base catalyst, deprotonating the water molecules when it attacks. When catalyzing substrates, Gln250 and His252 interact with catalytic residues, and participate in substrate binding to affect catalysis (Figure 5).

**Figure 5.**
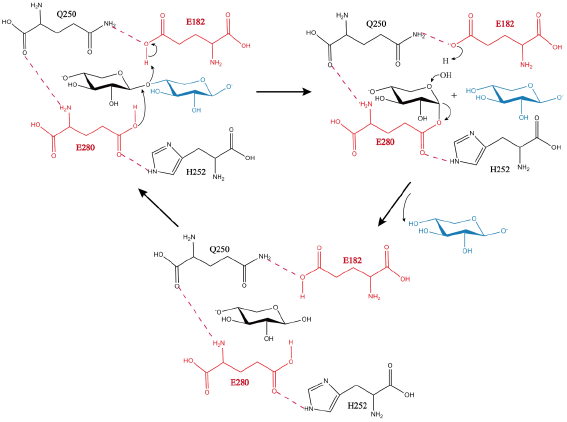
Proposed endo-1,4-β-xylanase catalytic mechanism for XynA. E280 acts as a nucleophile, and E182 acts as an acid/base catalyst. Q250 and H252 will interact with catalytic residues. The blue sugar represents the sugar that has been hydrolyzed. The pink dotted line represents hydrogen bond interaction.

XynA is a heat-stable and acid-stable xylanase in the GH10 family, and Celluclast 1.5L can synergistically hydrolyze pretreated corn stover. These properties indicate that it may be effective in food, animal feed and biofuel production. Analysis of the XynA crystal structure can provide a basis for its transformation, making it more suitable for commercial applications such as biofuel production.

## Materials and Methods

### Expression and purification of the wild-type and mutant XynA proteins

*E. coli* strains DH5α and BL21-CodonPlus (DE3) were purchased from Shanghai Veidi Biotechnology Co, Ltd. DH5α was used to construct the expression vector pET28a-XynA (Merck, Germany), and BL21-CodonPlus (DE3) was used to express XynA. The GenBank of XynA is QCO69162. The target protein was preceded by a 6xHis tag. The recombinant plasmid pET28a-XynA was transformed into BL21-Codonplus (DE3) competent cells, and then a single colony was selected and inoculated into 100 mL of LB medium containing 30 μg/mL kanamycin (Sangon Biotech, Shanghai, China) and 32 μg/mL chloramphenicol (Sangon Biotech, Shanghai, China) and cultured in a 37 °C constant temperature shaking table at 180 rpm for 10 hours. When the OD_600_ reached 0.6-0.8, the *E. coli* had reached the logarithmic growth stage. Protein expression was then induced with 0.2 mM isopropyl-β-D-thiogalactopyranoside (IPTG, Sangon Biotech, Shanghai, China) at 25 °C on a constant temperature shaker for 12 hours. Finally, the samples were collected by centrifugation at 4000 rpm for 20 minutes at 4 °C and stored at -80 °C until use.

The collected *E. coli* was suspended in buffer A containing 40 mM Tris-HCl at pH 8.0 with 250 mM NaCl and 10 mM imidazole at 4 °C, and then 1 mM β-mercaptoethanol (Amresco, Guangzhou, China) and 1 mM PMSF (Sigma, Beijing, China) were added. The resuspended *E. coli* cells were passed through a low-temperature ultrahigh-pressure cell disrupter at 1200 psi, and the obtained lysate was centrifuged at 14000 rpm for 60 minutes (4 °C). After centrifugation, portions of the supernatant and the precipitate were retained (coomassie brilliant blue: running glue samples) (34). The supernatant was divided into two parts: one part was directly combined with nickel-nitrilotriacetic acid (Ni-NTA) affinity resin (Qiagen, China); the other part was heated in a 65 °C water bath for 60 minutes and then centrifuged at 14000 rpm for 60 minutes. After the second centrifugation, a part of the supernatant and the precipitate were retained, and the supernatant was combined with Ni-NTA. The two supernatants above were combined with Ni-NTA for 45-60 minutes. The hybrid protein was removed by using buffer A, and the target proteins were eluted by using buffer B (40 mM Tris-HCl at pH 8.0 with 250 mM NaCl, 250 mM imidazole at 4 °C).XynA was purified in two ways, one of which was heating in a 65 °C water bath for 60 minutes and the second was using a Hi Trap QHP column (1 mL GE Healthcare, United States). When loading, the protein is bound to the column. The conditions are then changed, and the bound components elute separately. Purification was carried out on a 1 mL QHP column with a NaCl gradient, and the protein solution was injected into the equilibrium column at a flow rate of 0.5 mL/ minute. A linear NaCl gradient elution of 50 to 250 mM was used, and 2 mL fractions of the eluate were collected over the entire gradient length.

The state of XynA in solution was analyzed by a Superdex 200 (10/300) increase column (GE Healthcare, United States). First, the chromatographic column was balanced with a buffer containing 150 mM NaCl, 40 mM Tris-HCl at pH 8.0 and 1 mM dithiothreitol (DTT, Sangon Biotech, Shanghai, China). Second, XynA loading. Finally, the column was eluted with buffer solution, and 2 mL fractions of eluate was collected.

### Protein crystallization, data collection and structure determination

The home-grown crystals were initially screened manually using a laboratory-made kit at room temperature using a 96-well plate with diffused vapor diffusion. A 0.8 μL aliquot of XynA protein solution (8.0 mg/mL, 20 mM Tris-HCl pH 8.0, 287 mM NaCl) was mixed with a 0.8 μL reservoir solution and equilibrated with a 100 μL reservoir solution. Twenty-four hours later, the initial XynA crystals were obtained in 28% PEG600 (w/v), 0.1 M CaCl_2_ and 0.1 M MES pH 6.0. The optimized crystallization conditions were 24% PEG600 (w/v), 0.2 M CaCl_2_, 0.1 M MES pH 6.0, crystal length reaches 2 mm. 29% PEG600 (w/v), 0.1 M CaCl_2_, 0.1 M MES pH 6.0 and 25% glycerol were used as refrigerating protectors. Individual crystals were installed on a nylon loop and immediately cooled with liquid nitrogen prior to data collection. The individual crystals are installed on a nylon loop and immediately cooled with liquid nitrogen. In Shanghai Synchrotron Radiation Facility (SSRF-BL19U1) diffraction data was collected through X-ray diffraction and finally processed using HKL2000 (35). We use PDB 1V0L as a model to perform molecular replacement to obtain the initial model. Based on the initial model, we use COOT to accurately modify the structure, and then use CCP4 for refinement. Finally, repeat the above steps until the structural requirements are met (36, 37). XynA finally obtained a 2.3Å resolution crystal structure. The crystallization conditions of the XynA E182A/E280A were the same as those of XynA, and the structure of the XynA E182A/E280A was obtained by MR with a resolution of 2.9 Å. Information about the protein crystal structure is provided in Table 1.

### Site-directed mutagenesis

We designed primer pairs with mutation points. The primers of E182A were used to cause the plasmid pET28a-XynA to undergo PCR. Then, the PCR product was digested with *Dpn*I (Thermo Scientific, China) for 1.5 hours in a water bath at 37 °C, and the product was heated at 80 °C for 5 minutes to denature. Finally, the product was transformed into competent DH5α cells. The mutants were transformed into BL21-CodonPlus (DE3) cells to express the protein. The mutant E280A needed to be mutated on the plasmid pET28a-XynA (E182A) of the constructed mutant E182A. XynA E182Q/E280Q had the same construction method as XynA E182A/E280A. The purification method for the mutant XynA E182A/E280A and XynA E182Q/E280Q were the same as that of XynA.

### Multiple residue sequence alignment

We downloaded the residue sequences of the GH10 family of xylanases such as CbXyn10C and SlXyn10A from the GenBank protein database. Then, we compared the residue sequences of XynA, the residue sequence of the GH10 family of monofunctional xylanases, and the residue sequence of the GH10 family of bifunctional xylanases using ClustalW software and the ESPript3 website.

### Protein structure conservation analysis

We output the XynA protein structure that we parsed in pdb format. Second, we opened the ConSurf Server domain based on the prediction of protein sequence similarity or structure surface important functional areas) and selected “Amino Acids” and “known protein structure”. Third, we upload the pdb file of XynA for automatic analysis. Finally, we download the output results from the website and used PyMOL software to prepare the images.

### Biolayer interferometry binding studies

The ForteBio Octet^®^ RED96 interaction analyzer (ForteBio, United States) was used to detect the binding of XynA to the oligosaccharide substrates (cellobiose, xylohexaose and cellohexaose) (Megazyme, Ireland) and xylan (BWX) (Shanghai, China) (38, 39). First, XynA was dialyzed against PBS. Second, the equivalent biotinylation kit (Biotinylation Kit, Jiangsu Bomeida Life Science Co., Ltd.) was added to bind XynA protein for 60 minutes for biotinylating. Finally, the unbound biotin was removed with a desalination column. Before the experiment, we wetted the biosensor (SSA, Shenzhen Baokeda Biotechnology Co., Ltd.) with PBS for 10 minutes. In the first step, we equilibrated the sensor with PBS for 2 minutes. In the second step, XynA was loaded onto the SSA sensor until it was saturated. In the third step, we equilibrated with PBS for 3 minutes. In the fourth step, diluted substrate solutions (12.5, 25, 50, 100, 200, and 250 μM) were sequentially added and dissociated from low concentration to high concentration, and each combination and dissociation time was 300 seconds. A sensor not bound to the biotinylated XynA was used as a baseline control for calibration. The whole process was carried out at room temperature, and all tests were repeated three times.

### Protein thermal shift assay

SYPRO Orange Protein Gel Stain 5000× (Thermo Scientific, United States) was diluted with protein buffer solution (20 mM Tris-HCl at pH 8.0 with 287 mM NaCl) to 200×. A 30 μg portion of protein (5.0 mg/mL) was mixed with BWX and incubated on ice for 60 minutes, then centrifuged at 14000 rpm/ minutes for 10 minutes. After centrifugation, the protein was evenly mixed with 2.5 μL 200× SYPRO Orange, and the total volume was 20 μL. The mixture was heated from 25 °C to 99 °C at a rate of 0.2 °C/minute, and each sample was prepared in triplicate. An Applied Biosystems real-time PCR instrument (7500 Fast, Thermo Scientific, United States) was used to determine the protein denaturation temperature (exposure temperature of hydrophobic amino acid residues), and the results were generated using Origin software (40).

### Circular dichroism analysis

We use a circular dichroism spectrometer (Applied Photophysics Limited, United Kingdom) to measure the circular dichroism of proteins. We placed the sample in a 1 mm quartz colorimetric plate at room temperature (25 °C) and scanned the protein solution (0.4 mg/mL protein dissolved in 20 mM Tris-HCl at pH 8.0 buffer) within a range of 190 nm to 300 nm. We will take three consecutive scans and then average them to obtain the spectrum for each wavelength. The content of protein secondary structures was calculated by CDpro software using values from 190 nM to 300 nM.

### Molecular docking

First, the PDB files of XynA, xylohexaose and cellohexaose were prepared using the Protein Preparation Wizard of Maestro (Maestro11.9, Schrödinger, LLC, New York, 2019) to prepare the model structure for docking (41, 42). Then, the appropriate docking pocket was selected, and the molecular docking was performed. The docking scores of each ligand were determined with the best posture on Maestro. The criterion for hydrogen bond judgment is that the distance between the acceptor and donor atoms should be less than 3 Å. Output the docking file through Maestro, and finally use PyMOL software to make frigure.

### Determination of the enzyme activity by DNS

The DNS method was used to detect the xylanase activity of the XynA and XynA E182A/E280A proteins. In this study, BWX (Lot# X4252, ≥ 90% purity) was used as the substrate, and xylose was used as the standard. The reducing sugars released from BWX were measured by DNS reagent to estimate the xylanase activity. The hydrolyzed product oligosaccharides (reducing sugars) undergo redox reactions with 3,5-dinitrosalicylic acid (DNS) to produce 3-amino-5-nitrosalicylic acid (brown red). First, we took the appropriate concentration (25, 250, 500, 1000 nM) of the protein and 0.5% (w/v) of BWX and mixed them evenly in 100 mM sodium citrate at pH 6.5, and the total volume was 50 μL. Second, the reaction system was incubated at the optimal temperature of 65 °C for 10 minutes. Third, the DNS reagent was added to the reaction system, and the mixture was incubated at 95 °C for 5 minutes. Finally, the reaction system was cooled in an ice bath to room temperature and diluted with 100 mM sodium citrate buffer solution at pH 6.5, and the absorbance was determined at 540 nm with an enzyme label analyzer.

### Determination of the enzyme activity by the pHBAH method

The *p*HBAH method was used to detect the xylanase activities and cellulase activities of the different XynA mutants relative to XynA (43). First, XynA and its mutants were diluted appropriately. Second, the proteins were separately incubated with xylan substrates (WAX and BWX) and the cellulose substrate (barley β-glucan) in 50 mM Na_2_HPO_4_-NaH_2_PO_4_ buffer (pH 6.0) at 65 °C. The reaction time was 5 minutes for xylan substrates and 30 minutes for the cellulose substrate. Finally, the *p*-hydroxybenzoyl hydrazine (redox reaction with reducing sugar) reagent was added to determine the released reducing sugar, and the resulting *p*NP (*p*-nitrophenol) was measured by spectrophotometry at 410 nm.

## Data availability

The atomic coordinates and structure factors of xylanase XynA structure has been uploaded to the Protein Data Bank. All remaining data are included in the article.

## Acknowledgements

This work was supported by the following grants: the Youth Scientific Research Training Project of GZUCM (No. 2019QNPY07), the National Natural Science Foundation of China (No. 31700625), Guangdong Key Laboratory for Translational Cancer Research of Chinese Medicine (No. 2018B030322011), the National Natural Science Foundation of China (31802099), the Pearl River Talent Recruitment Program of Guangdong Province (2017GC010372), the Natural Science of Foundation of Guangdong Province, China (2018A030310497), the Research Program for Young Innovative Talents in Higher Education Institutions of Guangdong Province (2017KQNCX037) and Shanghai Synchrotron Radiation Facility.

## Author contributions

Caiyan Wang, Kui Wang and Zhongqiu Liu: Conceptualization, Methodology.

Wei Xie: Data curation, Writing-Original draft preparation.

Wei Xie, Qi Yu, Yun Liu, Ruoting Cao, Ruiqing Zhang, Sidi Wang and Ruoting Zhan: Software.

Wei Xie, Qi Yu, Caiyan Wang and Kui Wang: Visualization, Investigation.

Caiyan Wang, Kui Wang and Zhongqiu Liu: Writing-Reviewing and Editing.

